# ProtPen combines sequence- and structure-based approaches to facilitate protein function predictions on a proteome-wide scale

**DOI:** 10.64898/2026.07.11.737882

**Authors:** Diya Mathai, Stefan Schulze

**Affiliations:** Gosnell School of Life Sciences, Rochester Institute of Technology, Rochester, NY 14623, USA

**Keywords:** protein function annotation, proteomics, microbiology, protein structure, orthology

## Abstract

Proteins of unknown function represent a significant gap in our understanding of biological processes, encompassing large portions of the proteomes of many organisms, especially prokaryotes. Addressing this gap is critical to understanding the biology and pathogenicity of such organisms. We introduce ProtPen, an open-source pipeline that facilitates protein function prediction by combining eggNOG-mapper for sequence-based annotation with Foldseek for rapid structural similarity searches using AlphaFold-predicted protein structures. Annotation results from both tools are merged and enriched with UniProt metadata to produce a comprehensive output suitable for downstream analysis. The pipeline requires only a FASTA input file with UniProt identifiers, and is designed to analyze datasets on the scale of whole proteomes. Benchmarking on a curated dataset of well-characterized *Pseudomonas aeruginosa* proteins demonstrated an annotation accuracy of >90%, and highlighted the complementarity of sequence- and structure-based methods. Further evaluation of ProtPen included its application to biologically relevant datasets, comprising proteins of unknown function that exhibited significant differential abundances in a proteomics dataset of *P. aeruginosa*, and uncharacterized glycoproteins from *Haloferax volcanii*. ProtPen is readily extensible to incorporate additional protein function prediction tools. In summary, this pipeline facilitates the systemwide annotation of proteins of unknown function from proteomic datasets and whole proteomes.

**For Table of Contents Only:** 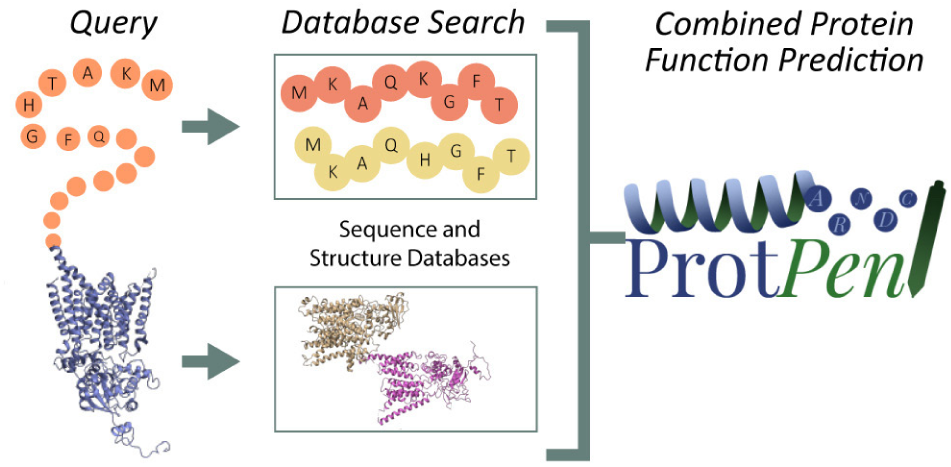

## Introduction

The advancement of next-generation sequencing (NGS) technologies and the rise of metagenomics has led to a large increase in the number of sequenced whole genomes across all domains of life. Public databases now host hundreds of complete and draft genomes, spanning from well-characterized organisms to rare and understudied taxa ^1,2^. As these genomes are annotated, they yield corresponding theoretical proteomes containing comprehensive catalogs of predicted proteins derived from open reading frames and gene prediction algorithms ^3^. However, a significant portion of identified proteins remain poorly annotated or entirely uncharacterized, limiting biological interpretation of experimental datasets, especially for system-wide analyses like proteomics and transcriptomics. This challenge is particularly acute in prokaryotes, where studies estimate that up to 30–50% of proteins in bacterial proteomes lack functional annotation ^4^. In the extremophile archaeon *Haloferax volcanii*, for example, the percentage of proteins with hypothetical, probable, or unknown function is 29.5%^5,6^. Even for well-characterized model organisms like *Pseudomonas aeruginosa*, a member of the ESKAPE (*Enterococcus faecium*, *Staphylococcus aureus*, *Klebsiella pneumoniae*, *Acinetobacter baumannii*, *P. aeruginosa*) group of pathogens, 9% of the annotated genome consists of uncharacterized proteins ^7^.

Along with advances in genome sequencing, transcriptomics and proteomics have been used to gain insights into the roles of various proteins. However, while comparative, quantitative studies regularly show differential abundances for previously uncharacterized proteins, indicating their biological relevance, they can often only point to broad cellular processes that these proteins are involved in; determining their specific functions requires time- and resource-intensive follow-up studies. These constraints not only affect the ability to functionally annotate proteins on a large scale, but in turn also limit the interpretability of system-wide analyses. Therefore, scalable, bioinformatic annotation frameworks are needed that can leverage available sequence and structure information to enhance functional annotations, particularly for proteins that fall outside well-studied orthologous groups.

Traditional approaches for protein function annotation rely on sequence homology, where functional information is inferred by aligning a query structure to existing proteins with known functions. Tools such as BLAST ^8^ perform pairwise sequence alignments to detect regions of similarity, while profile Hidden Markov Model (HMM) tools like HMMER ^9^ capture more distant homologs by using probabilistic models trained on protein families. These methods perform well when close homologs are available and characterized, but quickly lose accuracy as sequence identity decreases. To improve annotation coverage, orthology-based frameworks such as eggNOG-mapper ^10^ utilize precomputed groups of evolutionarily related proteins, known as orthologous groups (OGs), to assign function across a broader taxonomic range. eggNOG-mapper combines fast sequence alignment via DIAMOND ^11^ or MMseqs2 ^12^ with curated annotations from the eggNOG database ^13^. This approach allows for the transfer of functional information from well-annotated genes to those in less-characterized organisms, including GO terms, KEGG pathways, enzyme classifications, and protein domains. Other sequence-based methods rely more heavily on GO terms to predict functions. Recently, this approach includes various machine learning-based tools, such as DeepGO, that have been trained with vast datasets of sequences and corresponding GO terms^14^. However, especially in prokaryotes, with the enormous functional diversity of their proteomes, GO terms may not capture the unique functions of many proteins, limiting their annotations to higher hierarchical levels^15^. Overall, sequence-based methods often struggle to provide informative predictions when a query protein lacks homologs with known function.

Leveraging the three-dimensional (3D) structure of proteins provides an alternative for sequence-based predictive analyses. Structural features, such as active sites, binding pockets, and interaction interfaces arise from the spatial arrangement of amino acids that are not necessarily neighboring each other, unlike their linear sequence. Furthermore, similar structures can be formed despite a lack of sequence conservation. Therefore, structure comparisons allow for the detection of functional relationships that may not be evident from sequence similarity alone.

With the advance of high-quality structure predictions since the development of AlphaFold2 ^16^, structure-based search tools can compare predicted protein structures not only against experimentally determined entries in the Protein Data Bank (PDB) ^17^, but also against predictions in the AlphaFold Protein Structure Database^18^. Various tools have emerged to perform structure comparisons. Dali ^19^, with its standalone application DaliLite.v5, facilitates protein structural alignment and database searches by employing distance matrix alignment for optimal pairwise matching. Focusing on structure-centric protein function prediction, DeepFRI ^20^ employs a Graph Convolutional Network, using sequence attributes from a protein language model as well as protein structures. Furthermore, Foldseek ^21^ offers a scalable and computationally efficient approach to structural comparison. It converts 3D protein structures into simplified, sequence-like representations that facilitate rapid alignment using methods analogous to traditional sequence search algorithms. This enables Foldseek to perform large-scale structure-based searches against databases such as AlphaFoldDB or the PDB, identifying remote structural homologs at a speed and scale previously unattainable with tools like Dali. Several recently published pipelines leverage protein structural information to extend functional annotation beyond the reach of sequence homology alone. MorF (MorphologFinder) proposes a workflow of openly available tools for functional annotation via structural similarity, demonstrating that structure-based approaches can annotate an additional 50% of a proteome beyond standard sequence-based methods, with over 90% accuracy on proteins with known homology^22^. While MorF presents a proof-of-concept for a sequence-structure combined workflow, it is designed as a loosely coupled workflow rather than a packaged pipeline, requiring users to configure and run individual tools independently, which limits reproducibility and portability across computing environments. Phold further illustrates the strength of combining structural and sequential protein information to inform function, combining the ProstT5 protein language model with Foldseek to search against a curated database of over 1.36 million predicted phage protein structures, outperforming sequence-based homology approaches in annotation sensitivity while maintaining speed and scalability, but remains scoped to phage genomes.^23,24^

To benefit from the advantages of both sequence- and structure-based methods, while focussing on the analysis of whole proteomes from any organism, we developed ProtPen to assist in the challenging annotation of proteins of unknown function. ProtPen is a modular, Python-based pipeline that integrates sequence-based annotations from eggNOG-mapper with structure-based searches using Foldseek. ProtPen automates protein structure retrieval via the AlphaFoldDB API, performs high-throughput Foldseek searches, enriches structural hits with UniProt metadata, and merges these outputs with the various classifications provided by eggNOG-mapper into a comprehensive TSV file. The pipeline accepts FASTA input with UniProt identifiers, and supports reproducible, large-scale execution on computational clusters. We benchmark ProtPen on a curated dataset of *P. aeruginosa* proteins, as well as on two experimental proteomics datasets, demonstrating its reliability, scalability, and usefulness for gaining insights into uncharacterized proteins with demonstrated biological relevance. The pipeline is designed for extensibility and can be readily applied to other organisms or datasets where proteins of unknown function remain a barrier to discovery.

## Methods

### Workflow Overview and Implementation

ProtPen was designed as a Python-based workflow that automates each step of a protein annotation pipeline, including sequence-based searches, retrieving AlphaFold predicted structures for structure-based searches, as well as enhancing and consolidating the results. Its modular design allows for the implementation and replacement of tools as new methods emerge, enabling continued adaptation to evolving bioinformatic resources.

The pipeline accepts as input a FASTA-formatted reference proteome in which protein headers include valid UniProt identifiers. A subset of proteins from this reference, e.g. derived from proteomic experiments or other specific datasets, can be used to define a list of query proteins for downstream structural annotation. ProtPen retrieves AlphaFold-predicted structures on demand for the query proteins. ProtPen requires a local installation of eggNOG-mapper and Foldseek, the latter of which comes with a pre-indexed Foldseek database. The full workflow consists of six stages: (1) eggNOG-mapper annotation based on sequence comparisons, (2) retrieval of AlphaFold structures for query proteins, (3) Foldseek structural search against the PDB, (4) filtering and consolidation of Foldseek results, (5) enrichment/enhancement of Foldseek hits using UniProt metadata, and (6) merging of sequence- and structure-based annotation tables into a unified output (Fig. 1). Each of these steps consists of a Python script as detailed below, and the complete pipeline has been wrapped as a Simple Linux Utility for Resource Management (SLURM) workflow. The workflow also facilitates multiprocessing to run eggNOG-mapper and Foldseek in parallel, and to process structure retrieval and result processing efficiently.

**Figure 1.**
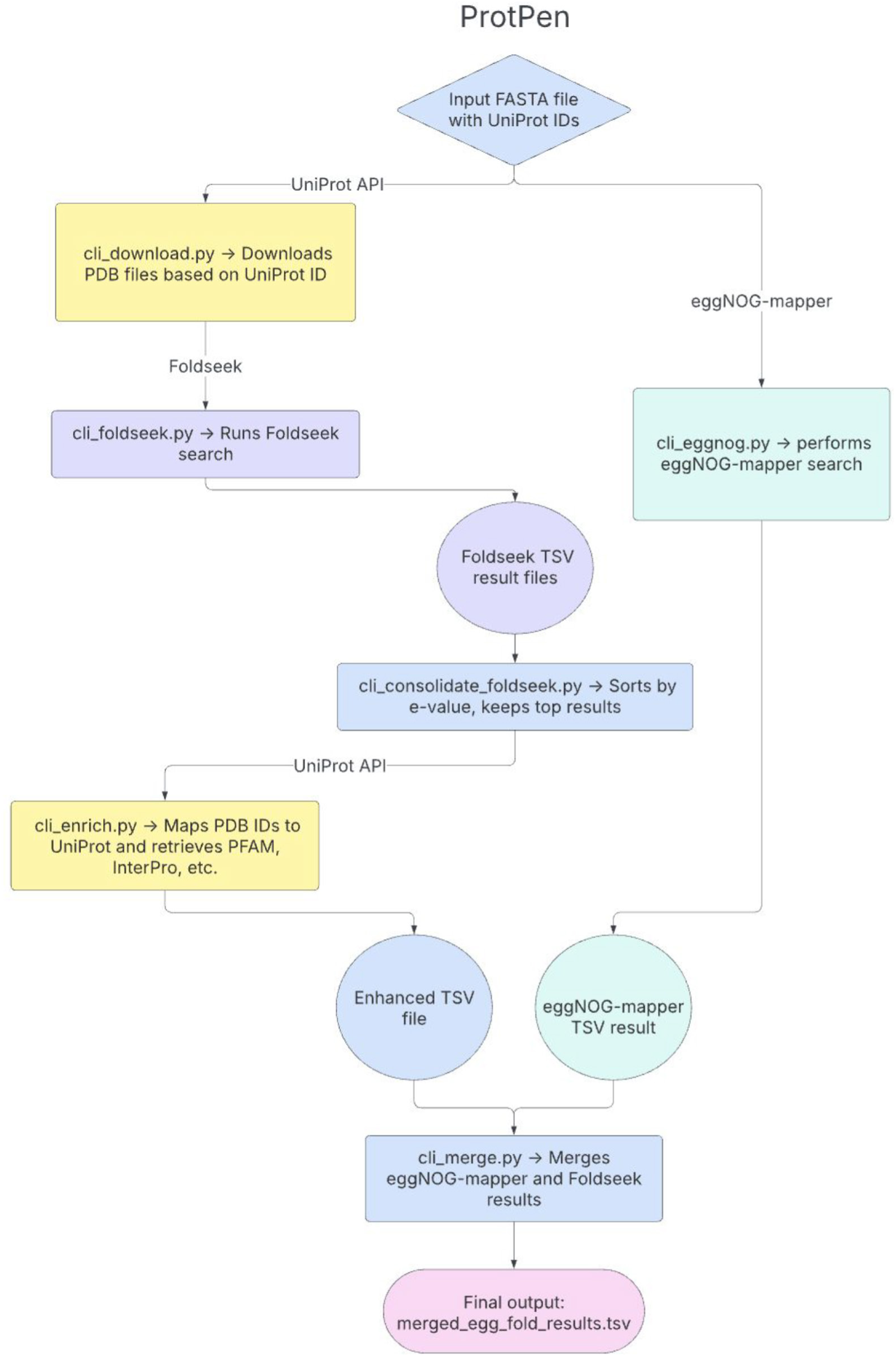
ProtPen workflow schematic, outlining the pipeline for protein function prediction using sequence- and structure-based approaches. The workflow begins with an input file: a reference proteome FASTA file containing protein sequences with UniProt IDs (blue). The FASTA file is passed to eggNOG-mapper (green) for sequence-based searches to generate functional annotations, producing a TSV file with gene ontology terms, KEGG pathways, and other annotations. AlphaFold PDB structures are retrieved (yellow) and then Foldseek analysis (purple) is performed, where the PDB structures are used in a structural similarity search, generating TSV result files. After these initial steps, the results from both eggNOG-mapper and Foldseek are filtered, retaining only entries corresponding to the query proteins. The filtered Foldseek results are then enhanced by retrieving additional protein domain information, including SUPFAM and InterPro annotations (yellow) based on their UniProt entries. Finally, in the merging step (blue), the enriched Foldseek results are combined with the eggNOG-mapper functional annotations, producing a final integrated output (pink).

During pipeline development, in addition to eggNOG-mapper and Foldseek, we evaluated several tools that offer sequence- or structure-based functional annotation capabilities, including Mantis ^25^, DeepFRI ^20^, and DALI ^19^. However, integrating these tools into a scalable, automated workflow proved difficult due to issues such as dependency conflicts, installation complexity, high resource demands, or lack of compatibility with computational cluster environments.

### Sequence-Based Annotation Using eggNOG-mapper

ProtPen uses `cli_eggnog.py` to annotate the reference proteome using eggNOG-mapper (version 2.1.12) ^10^, which assigns orthologous groups, Gene Ontology (GO) terms, enzyme codes, and KEGG pathway annotations based on homology. MMseq2 (version 113e3212c137d026e297c7540e1fcd039f6812b1) ^12^ is used as the default alignment method due to its speed and scalability. The resulting annotation is saved as a tab-delimited file, which is later filtered to retain only query proteins of interest. Default parameters for eggNOG-mapper have been used for all datasets described here. These parameters include searching against the eggNOG v5 database^13^, an e-value threshold of 0.001, --dbtype 1, --start-sens 3, --sens-steps 3, and -s 7; the --tax_scope option was not applied.

### Retrieval of AlphaFold Structures

Predicted protein structure models for query proteins are retrieved using `cli_download.py`. This script extracts UniProt IDs from the input FASTA file and queries the AlphaFold Protein Structure Database via the UniProt REST API ^16,26^. Available PDB files are downloaded and stored in a local directory for use in downstream Foldseek searches.

### Structural Similarity Searches Using Foldseek

Using Foldseek (version 9-427df8a) via cli_foldseek.py, each downloaded structure is compared to a database of known protein structures, using the easy-search method with default parameters ^21^. These default parameters aim to provide a balance between speed, sensitivity, and reliability, e.g. using -s 9.5 but no iterative or exhaustive search, and applying an e-value threshold of 0.001. We perform Foldseek searches against the PDB to prioritize experimentally verified structures; however, Foldseek also supports multiple AlphaFold-based databases. The ProtPen implementation supports batch processing, optional temporary directories, and multi-threaded execution, enabling efficient use of high-performance computational resources. Foldseek outputs one result file per query protein in TSV format, containing alignment statistics and structural similarity scores.

### Filtering and Consolidation of Foldseek Results

The Foldseek output is subsequently filtered and consolidated using `cli_consolidate_foldseek.py.` This module applies optional thresholds for e-value and top-N hit selection (e.g., top five matches per query). The filtered and ranked results are written to a single TSV file, simplifying downstream enrichment and integration.

### Enrichment with UniProt Metadata

To improve the interpretability of structural annotations, Foldseek results are enriched using `cli_enrich.py.` Each structural match is mapped back to a UniProt entry via the RCSB PDB GraphQL-based API. The UniProt API is then used to access metadata including protein names, InterPro domains, and SUPFAM classifications, which are appended to the consolidated results. This produces an enhanced TSV file that combines structural similarity information with corresponding functional information.

### Merging Annotation Sources

The final stage of the pipeline is performed using `cli_merge.py,` which integrates the enhanced Foldseek output with the filtered eggNOG-mapper annotations. Entries are matched by UniProt identifier, and the merged result includes columns from both tools. Multiple Foldseek matches per protein are retained and represented using a custom delimiter (||) for downstream parsing. The resulting merged TSV file provides a comprehensive annotation table for all query proteins, combining sequence homology, structural similarity, and functional information.

### Command-Line Interface and Execution Environment

ProtPen is implemented as a suite of modular command-line wrappers designed for execution on high-performance clusters with SLURM job scheduling. Each CLI script corresponds to one pipeline stage, and a full SLURM job script `(run_pipeline.sh`) demonstrates their sequential execution. However, SLURM job scheduling is not required, and each script can be executed individually as described in the documentation. Additional scripts handle result consolidation, metadata enrichment, and final merging of annotation outputs.

### Retrieving Datasets to Test ProtPen’s Functionality

Three types of datasets were used in this study. A control dataset of well-characterized *Pseudomonas aeruginosa* PAO1 proteins was curated from UniProtKB by selecting reviewed entries with experimental evidence at the protein level. These sequences were downloaded in FASTA format and used to benchmark pipeline performance. Complete reference proteomes were downloaded from UniProtKB for *P. aeruginosa* PAO1, *Neisseria gonorrhoeae* FA614 , *Haloferax volcanii* DS2, and *Pyrococcus furiosus* ATCC 43587. These proteomes served as inputs for sequence-based annotation, AlphaFold structure retrieval, and Foldseek searches. Two experimental datasets were analyzed: PRIDE dataset PXD044807 and the *H. volcanii* DS2 glycoproteome^27,28^. Protein identifiers from both studies were mapped to UniProtKB accessions and used as query sets within their respective reference proteomes.

### Assessment of ProtPen’s Results

ProtPen’s results were assessed through following three different approaches:

A. For proteins with known function (e.g. in the control dataset and the experimental datasets), outputs from eggNOG-mapper and Foldseek were manually inspected, compared against the known function, and assigned to one of three categories: “Matching”, indicating a specific annotation consistent with the known function; “Needs Review”, used for annotations that were partial, unspecific, or ambiguous with respect to the known function; and “Conflicting”, denoting a clear disagreement with the known function.
B. For uncharacterized proteins, basic annotation quality was assessed programmatically using a custom Python script. Each annotation was placed into one of three categories: Missing”, if the annotation field was absent, blank, or contained only a placeholder such as “-”; “Unclear”, when all annotation terms were uninformative, defined as containing the keywords “uncharacterized”, “hypothetical”, “DUF”, or “unknown”; and “Present”, when at least one informative term was present. Where multiple annotation terms were present in a single field, these were parsed by splitting on the “||” delimiter, and a score of “Present” was assigned if at least one individual term was informative.
C. For the comparison of results from eggNOG-mapper and Foldseek, each tool’s results were evaluated manually and classified as “Agree”, if both tools returned compatible

annotations; “Unclear”, when the comparison was ambiguous or required further review; or “Disagree”, if the two tools returned incompatible or contradictory annotations.

The assigned categories for each protein can be found in the corresponding Supplemental Data files for each dataset.

### Data and Code Availability

ProtPen is freely available under a GNU General Public License v3.0 at https://github.com/ProtPen/ProtPen where documentation and example scripts, as well as an explanation of the extensibility of ProtPen together with guidelines for community contributions, are provided. ProtPen can be installed via PyPI.

## Results and Discussion

### ProtPen integrates potential protein annotations from sequence- and structure-based approaches

ProtPen is a modular, Python-based pipeline designed to automate the prediction of protein functions using both sequence- and structure-based methods. The pipeline integrates orthology-based sequence annotation via eggNOG-mapper, structure-based similarity searches using Foldseek, and retrieval of additional information through the UniProt API. Each component can be executed individually or as part of a unified, SLURM-compatible job script. ProtPen was developed for deployment on high-performance computing (HPC) systems and is suitable for processing whole proteomes or large proteomic datasets.

### Assessing performance via a defined control dataset

To assess the performance of ProtPen, especially for prokaryotic proteomes, we created a curated control dataset composed of 115 *Pseudomonas aeruginosa* proteins with experimentally verified functions. This dataset was constructed by selecting proteins with protein-level evidence in UniProt and confirmed structural information in the PDB. Functional classes were selected to reflect a diverse set of biological roles, including metabolic enzymes, membrane transporters, lectins, pili-associated proteins, and regulatory components (Fig. 2A).

**Figure 2.**
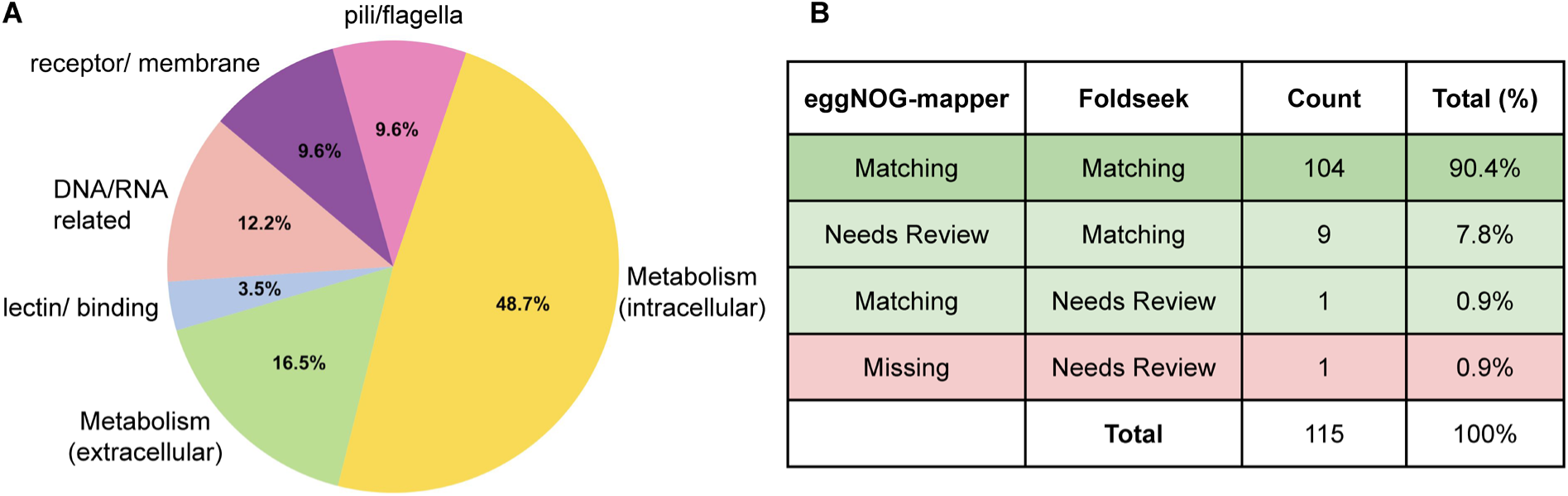
Performance of ProtPen on a control dataset of proteins with known functions. **(A)** Overview of protein categories in the control dataset from *Pseudomonas aeruginosa* PAO1. Categories comprised intracellular and extracellular metabolic proteins, pili/flagella-associated proteins, membrane-associated proteins and receptors, proteins that bind or synthesize nucleic acids, and lectins/carbohydrate-binding proteins. **(B)** Summary of annotation results for the control dataset. The results of the two tools currently implemented in ProtPen, eggNOG-mapper and Foldseek, were manually compared to the protein’s known function. “Matching” indicates specific annotations that are in line with the known function; “Needs Review” is used for partial/unspecific or ambiguous matches; “Conflicting” denotes clear disagreement from the known function, but was not observed for this control dataset (Supplemental Data 1).

After processing all protein sequences with ProtPen, the resulting outputs were compared with the known biological function for each protein and categorized as follows: (i), ProtPen results that clearly matched the known function, (ii), annotations that were ambiguous or too broad, requiring further manual review, and (iii), suggested annotations that conflicted with the known functional description (Supplemental Data 1). Due to the lack of a standardized nomenclature for protein descriptions, protein domains with related functions, and other functional descriptors, the proteins were assigned to each category through manual inspection of each protein’s functional description, as well as eggNOG-mapper and Foldseek results. Out of 115 proteins, 104 (90.4%) received annotations from both eggNOG-mapper and Foldseek that agreed with their known function (Fig. 2B). These results indicate a high degree of consistency between sequence- and structure-based approaches, and reliable annotations when the results from both tools provided similar functional information.

An additional 10 (8.7%) proteins showed matching results from one tool while requiring further review for results from the other. In these cases, annotations may be ambiguous or require more specific domain knowledge to assess their concrete functions. The more specific annotations from the second tool assist in determining likely functions, and they match the known functions of the characterized proteins, highlighting the benefits of combining multiple tools. Only 1 (0.9%) protein was missing annotations from eggNOG-mapper and the results from Foldseek did not clearly match the known function. A functional annotation for these proteins was therefore not possible.

Foldseek correctly annotated 113 proteins of our test dataset, compared to 105 by eggNOG-mapper. This difference was most apparent in proteins with low sequence similarity to known homologs, where structural alignment enabled functional inference not captured by sequence-based methods.

When evaluating ProtPen results, we consider results for which both eggNOG-mapper and Foldseek agree as most reliable. While all of the proteins with agreement between both tools matched the known protein function, suggesting a high accuracy, it should be noted that this control dataset represents only well-characterized proteins, which are thus well-represented in the employed databases. The accuracy of annotations for uncharacterized proteins cannot be evaluated with this approach, as the true function of these proteins is unknown. Similarly, while for none of the proteins in the control dataset, the obtained results clearly conflicted with the known functions of the proteins, indicating a low likelihood for false annotations, no conclusions about the false annotation rate for uncharacterized proteins can be drawn.

### Analyzing uncharacterized proteins of entire proteomes

To evaluate the scalability of ProtPen and its usefulness for different species, we applied the pipeline to the complete theoretical proteomes of *Pseudomonas aeruginosa, Neisseria gonorrhoeae, Haloferax volcanii*, and *Pyrococcus furiosus*. The runtime of ProtPen followed a linear correlation with the number of proteins in a proteome (Fig. 3A), and the median time for processing a protein was 5.1 seconds (Fig. 3B).

**Figure 3.**
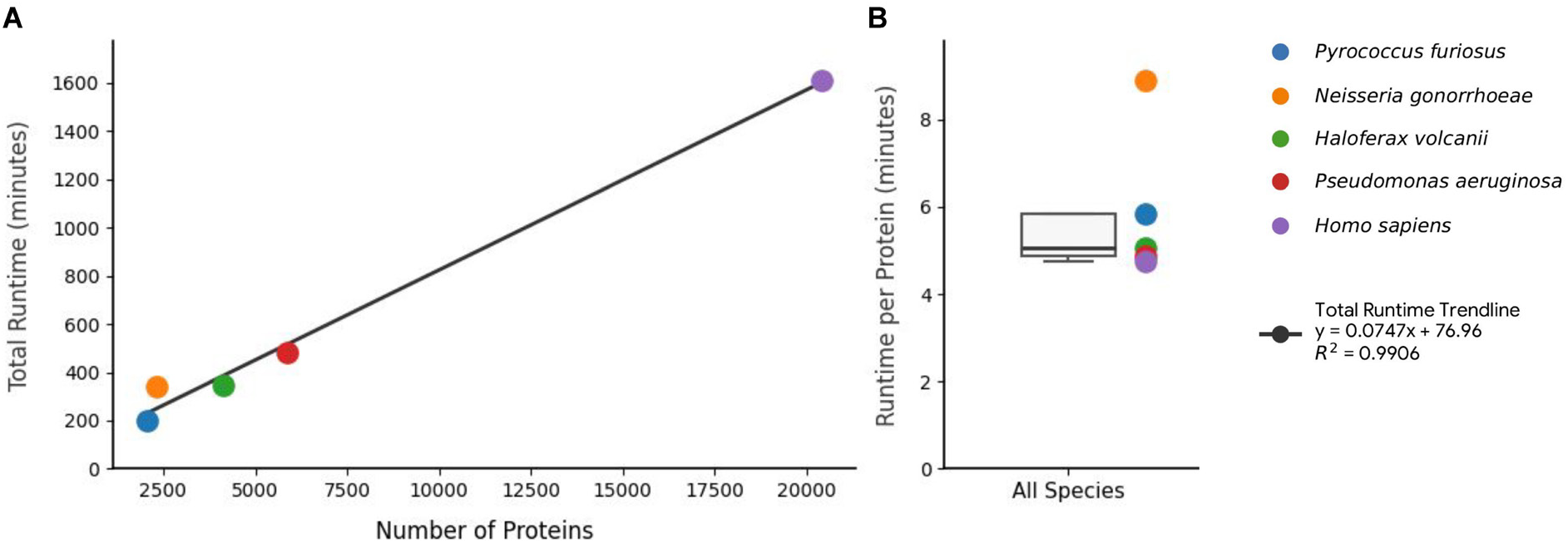
Runtime of ProtPen in relation to the proteome size. **(A)** The total runtime of ProtPen is depicted for each analyzed proteome in relation to the number of proteins in the respective proteome. In addition to the four analyzed prokaryotic proteomes, the proteome of *Homo sapiens* was included as a reference for comparatively large proteomes. The trendline represents the result of a linear regression. **(B)** The runtime per protein across all analyzed species is depicted as a boxplot with the line indicating the median (5.1 seconds) and the box representing the interquartile range. All analyses were performed using 16 CPU cores (Intel^®^ Xeon^®^ Gold 6150, 2.7 GHz) with a peak memory usage of 75 GB.

The species that we chose for our evaluations span both prokaryotic domains of life, Bacteria and Archaea. The Gram-negative bacterial pathogens *P. aeruginosa* and *N. gonorrhoeae* have been extensively studied due to their clinical relevance ^29,30^. Nevertheless, their proteomes contain a substantial portion of uncharacterized proteins. In *P. aeruginosa* PAO1, 9.0% of the proteome lacked functional annotation, while 11.2% of *N. gonorrhoeae* FA614 proteins were uncharacterized (Fig. 4). These results highlight the need for an improved characterization of bacterial proteomes.

**Figure 4.**
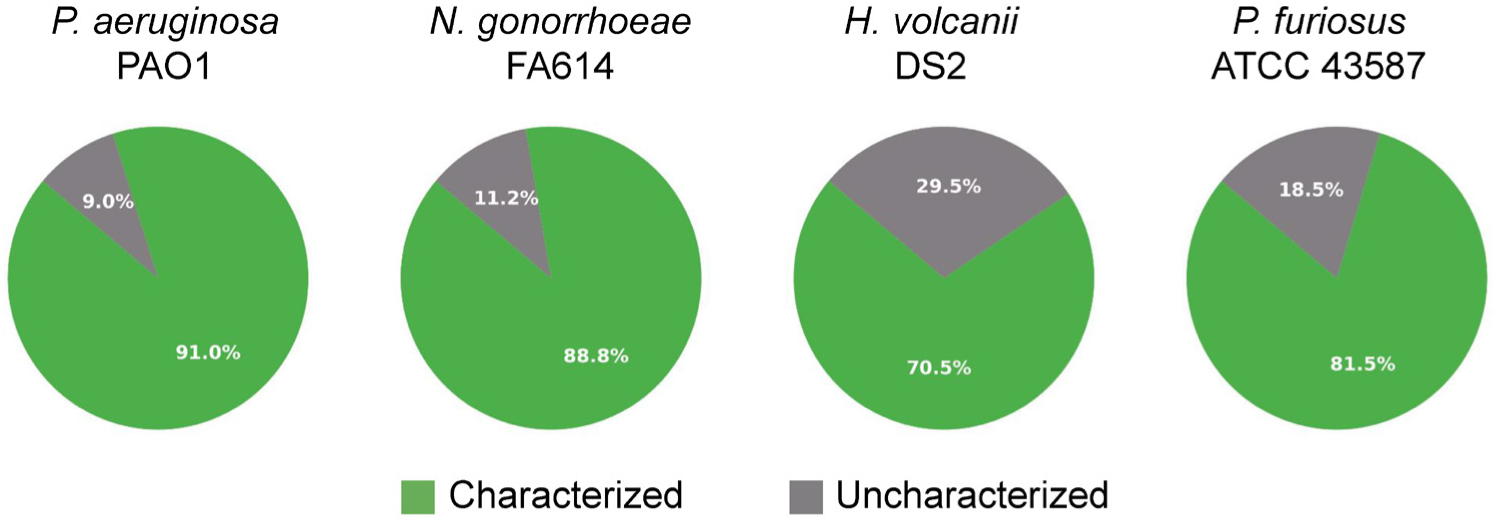
Proportion of characterized and uncharacterized proteins across four reference proteomes. Each pie chart displays the percentage of proteins annotated as characterized (green) or uncharacterized (gray) in the reference proteomes of *Pseudomonas aeruginosa* PAO1 (5563 proteins), *Neisseria gonorrhoeae* FA614 (2308 proteins), *Haloferax volcanii* DS2 (4186 proteins), and *Pyrococcus furiosus* ATCC 43587 (2044 proteins). Uncharacterized proteins include those labeled as hypothetical, probable, or of unknown function in the annotated genome. Percentages are calculated relative to the total number of proteins in each species’ proteome.

Similarly, while the two selected archaeal species, *H. volcanii* and *P. furiosus*, represent established model systems for halophilic and hyperthermophilic archaea, respectively, the fraction of their proteomes that corresponds to proteins of unknown function was even higher than for bacteria: 29.5% in *H. volcanii* DS2 and 18.5% in *P. furiosus* ATCC 43587^31,32^ (Fig. 4). This high percentage of uncharacterized proteins underscores the challenges of functional annotation in archaeal systems, which have been studied less intensively than many bacterial systems, and where divergence from well-studied organisms reduces the effectiveness of sequence-based homology searches.

Across all four species, ProtPen successfully processed the UniProt reference proteomes retrieving AlphaFold-predicted structures, running sequence-based annotation with eggNOG-mapper, and identifying structural homologs with Foldseek. For each organism, the merged output provided functional annotations for thousands of proteins (Supplemental Data 2). The extent of uncharacterized proteins varied widely among the species, but in each case ProtPen obtained potential functional annotations through at least one tool for a large proportion of uncharacterized proteins (Fig. 5A). While a direct comparison of tool performance across species is not feasible, due to the broad range of factors influencing results (e.g. proportion of uncharacterized proteins, phylogenetic distance, representation of phylegenetically related proteins in the employed databases), the presented results illustrate the applicability of ProtPen to species from different prokaryotic domains and with varying extensiveness of functional proteome annotation.

**Figure 5.**
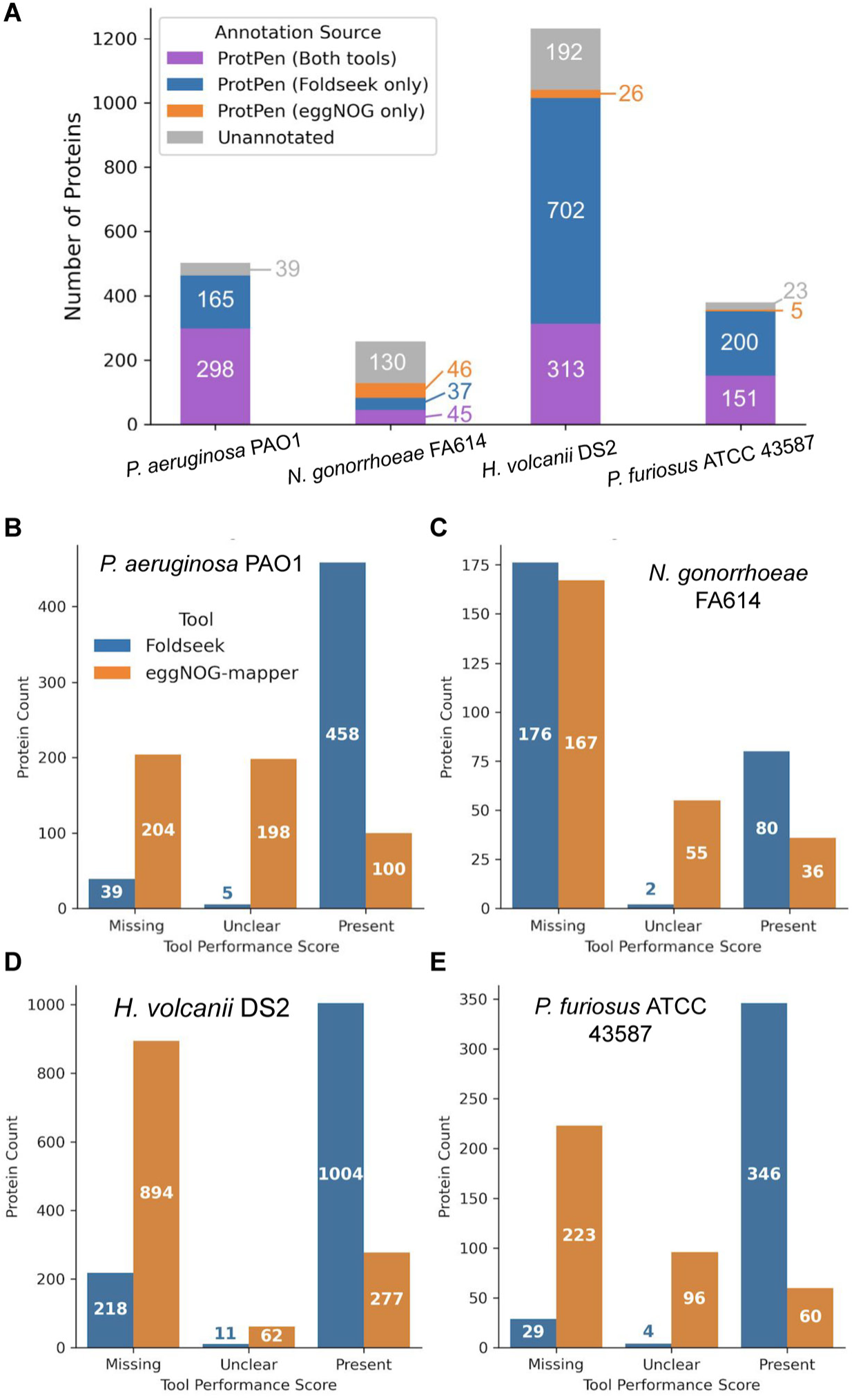
Annotation outcomes for uncharacterized proteins using ProtPen across four prokaryotic proteomes. **(A)** Stacked bar plot showing the annotation results for uncharacterized proteins from four species: *Pseudomonas aeruginosa* PAO1, *Neisseria gonorrhoeae* FA614, *Haloferax volcanii DS2*, and *Pyrococcus furiosus* ATCC 43587. Each bar shows proteins annotated by both tools (purple), by Foldseek only (blue), by eggNOG-mapper only (orange), or left unannotated (gray). **(B-E)** Bar plots depicting the number of uncharacterized proteins from each species annotated by Foldseek (blue) or eggNOG-mapper (orange), grouped by tool performance score: Missing (no information or blank entry), Unclear (vague or uninformative labels such as “hypothetical protein” or “DUF”) and Present (informative and specific annotations). Counts for each tool and species sum up to the total number of uncharacterized proteins from that species, i.e. each protein is counted separately for eggNOG-mapper and Foldseek. The classification of each protein for (A) and (B-E) can be found in Supplemental Data 2.

The *P. aeruginosa* PAO1 proteome contained 502 uncharacterized proteins, of which 298 (59.4%) were annotated by both eggNOG-mapper and Foldseek and 165 (32.9%) were annotated only by Foldseek, leaving the remainder unannotated (7.8%). The *H. volcanii* DS2 proteome contains a larger number of 1,233 uncharacterized proteins. Of these 25.4% and 56.9% was annotated by both tools and Foldseek only, respectively, and 26 (2.1%) proteins were annotated through eggNOG-mapper alone, while 192 (15.6%) remained unannotated.

The *N. gonorrhoeae* FA614 proteome had the smallest number of uncharacterized proteins (258), which likely contributed to a comparatively low annotation rate of ProtPen, with annotation from both tools for 45 (17.4%) proteins, from only Foldseek or eggNOG-mapper for 37 (14.3%) and 46 (17.8%), respectively, while 50.4% remained without potential function. For the 379 uncharacterized proteins of the *P. furiosus* ATCC 43587 proteome, ProtePen resulted in the annotation of 151 (39.8%) protein by both tools, 200 (52.8%) by Foldseek exclusively, 5 (1.3%) by eggNOG-mapper exclusively, and only 23 (6.1%) remaining without annotation (Fig. 5). It should be noted that the high annotation rates for *P. aeruginosa* PAO1 and *P. furiosus* ATCC 43587 proteomes likely reflect a strong representation of these model organisms, and their phylogenetic relatives, in structural and sequence databases.

Overall, these results demonstrate several consistent trends. First, dual annotation by both tools accounted for a large part of functional assignments (30-64% of proteins with at least one result returned by ProtPen), adding to the reliability of the predictions. Second, Foldseek contributed the vast majority of annotations when only one tool returned results, highlighting the value of structural similarity in uncovering protein function. Third, while ProtPen reduced the annotation gap substantially across all proteomes, a small fraction of proteins remained without sequence- or structure-based matches, reflecting the persistent challenge of functional characterization in prokaryotes.

To evaluate if sequence- and structure-based methods would result in a consensus annotation for uncharacterized proteins, results from both eggNOG-mapper and Foldseek for proteins of unknown function from *P. aeruginosa* PAO1 were compared (Table 1). For 91 uncharacterized proteins, both tools returned potential functional annotations that went beyond vague information such as “DUF”, “membrane” or “protein conserved in bacteria”. For the majority of uncharacterized protein (74.7%), both tools agreed on very similar annotations, indicating a high level of confidence for these results. The remaining proteins included 13 (14.3%) that needed further evaluation, and 10 (11%) for which the results from both tools were incompatible with each other (Supplemental Data 3). These results highlight the strength of ProtPen by combining sequence and structure-based functional information for uncharacterized proteins. Not only did the two implemented tools, eggNOG-mapper and Foldseek, agree on the potential functions for the majority of proteins of unknown function, their disagreement for a subset of proteins also allows us to recognize potential mismatches for each tool.

**Table 1.** Tool-to-tool comparison for uncharacterized *Pseudomonas aeruginosa* PAO1 proteins for which potential functional annotations were obtained by both eggNOG-mapper and Foldseek. . Agreement was assessed for each protein individually, with ‘Agree’ indicating compatible annotations between eggNOG-mapper and Foldseek, ‘Unclear’ corresponding to cases that require further review or showed ambiguous functions, and ‘Disagree’ referring to annotations between the tools that are incompatible or contradictory to each other (Supplemental Data 3). Only proteins with informative annotations by both tools (going beyond annotations such as “DUF”, “membrane” or “protein conserved in bacteria” were taken into account.

| Category | Proteins | (%) |
| --- | --- | --- |
| Agree | 68 | 74.7% |
| Unclear | 13 | 14.3% |
| Disagree | 10 | 11.0% |
| Total | 91 | 100% |

### Application of ProtPen to Experimental Datasets

To further establish ProtPen’s usefulness beyond theoretical proteomes, we extended its application to two experimental datasets that included uncharacterized proteins in their sets of biologically meaningful proteins: (1) the bacterial dataset PXD044807, which performed a quantitative proteomic analysis of *P. aeruginosa* PAO1 grown on different carbon substrates and reported multiple uncharacterized proteins with differential abundance, and (2) the *H. volcanii* glycoproteome, which was shown to include a large percentage of proteins of unknown function^27,28^.

### *Pseudomonas aeruginosa* Dataset PXD044807

The PXD044807 dataset, which compared protein abundances for *P. aeruginosa* grown on three fatty acid substrates relative to glucose, comprised 76 proteins that showed differential abundance^27^. Among these, 67 had known functions while 9 were previously uncharacterized. ProtPen confirmed or refined annotations for most of the characterized proteins, and successfully returned likely annotations for 3 of the uncharacterized proteins, while providing first insights into potential functions for 5 additional proteins (Table 2 and Table 3).

**Table 2.** Comparison of ProtPen-derived functional annotations with known functions for characterized, differentially abundant *P. aeruginosa* PAO1 proteins from dataset PXD044807. ^27^. “Matching” indicates specific annotations that are in line with the known function; “Needs Review” is used for partial/unspecific or ambiguous matches; “Conflicting” denotes clear disagreement from the known function (Supplemental Data 4).

| eggNOG-mapper | Foldseek | Proteins | (%) |
| --- | --- | --- | --- |
| Matching | Matching | 62 | 92.5% |
| Needs Review | Matching | 1 | 1.5% |
| Matching | Needs Review | 1 | 1.5% |
| Needs Review | Needs Review | 1 | 1.5% |
| Conflicting | Needs Review | 2 | 3.0% |
|  | **Total** | 67 | 100% |

**Table 3.** Assessment of ProtPen annotations for uncharacterized, differentially abundant *P. aeruginosa PAO1* proteins from dataset PXD044807. ^27^. Agreement between eggNOG-mapper and Foldseek was classified as Agree (compatible annotations), Unclear (ambiguous or requiring further review), or Disagree (incompatible or contradictory annotations). Only Foldseek and Only eggNOG-mapper indicate proteins for which informative annotations were provided by a single tool (Supplemental Data 4).

| Category | Proteins | (%) |
| --- | --- | --- |
| **Agree** | 3 | 33.3% |
| **Only Foldseek** | 3 | 33.3% |
| **Only eggNOG-mapper** | 1 | 11.1% |
| **Unclear** | 1 | 11.1% |
| **Disagree** | 1 | 11.1% |
| **Total** | 9 | 100% |

For 62 of the characterized proteins, the results from both Foldseek and eggNOG-mapper matched the known function of those proteins. 2 proteins had a matching result from one tool while the other tool required further investigation and 1 protein had ambiguous results from both tools. Additionally, for 2 characterized proteins the eggNOG-mapper results conflicted with the known function while the results from Foldseek were ambiguous. These results again indicate that ProtPen provides reliable annotations, especially when both the sequence- and structure-based tool agree (Table 2 and Supplemental Data 4).

Of the 9 uncharacterized proteins, 3 had annotations from both tools that aligned with each other, representing the most reliable category of ProtPen results (Table 3). All 3 annotations are in line with proteins that could be affected by metabolic changes due to the altered substrate availability: a potential long-chain fatty acid transport protein (Q9I456), a potential LysR-family transcriptional regulator (Q9HUP9), and a potential acetyl-coenzyme A synthetase (P28812) (Supplemental Data 4). While the annotations for 3 additional proteins were ambiguous for both tools, and 4 further proteins returned results only for one tool each, these results could serve as a starting point for more detailed reviews of the sequences and structures of the corresponding proteins. Only one uncharacterized protein had conflicting results from Foldseek and eggNOG-mapper.

Overall, ProtPen yielded consistent and complementary annotations for the vast majority of proteins with known functions in this dataset. In addition, the ProtPen analysis resulted in multiple functional predictions for previously uncharacterized proteins that showed differential abundance in the presence of varying fatty acid substrates, indicating their biological importance. These results underscore the reliability and value of ProtPen’s approach to combine sequence- and structure-based methods.

### *Haloferax volcanii* glycoproteins

To assess ProtPen’s utility in archaeal species, we analyzed a dataset of *H. volcanii N-*glycoproteins, encompassing 32 uncharacterized and 20 characterized proteins (Table 4 and Table 5). Similar to the other datasets described above, results for the vast majority of previously characterized glycoproteins matched the known function of the glycoproteins (Supplemental Data 5). Notably, no annotations were returned that conflicted with the known protein functions (Table 4).

**Table 4.** Comparison of ProtPen-derived functional annotations with known functions for characterized *N*-glycoproteins identified in *H. volcanii*^28^. “Matching” indicates specific annotations that are in line with the known function; “Needs Review” is used for partial/unspecific or ambiguous matches; “Conflicting” denotes clear disagreement from the known function (Supplemental Data 5).

| eggNOG-mapper | Foldseek | Proteins | (%) |
| --- | --- | --- | --- |
| Matching | Matching | 15 | 75% |
| Needs Review | Matching | 3 | 15% |
| Matching | Needs Review | 1 | 5% |
| Needs Review | Needs Review | 1 | 5% |
|  | **Total** | 20 | 100% |

**Table 5.** Assessment of ProtPen annotations for uncharacterized *N*-glycoproteins identified in *H. volcanii*^28^. Agreement between eggNOG-mapper and Foldseek was classified as Agree (compatible annotations), Unclear (ambiguous or requiring further review), or Disagree (incompatible or contradictory annotations). Only Foldseek and Only eggNOG-mapper indicate proteins for which informative annotations were provided by a single tool. Neither tool returned informative results for ‘Missing’ proteins (Supplemental Data 5).

| Category | Uncharacterized Proteins | (%) |
| --- | --- | --- |
| Agree | 3 | 9.4% |
| Only Foldseek | 15 | 46.9% |
| Only eggNOG-mapper | 1 | 3.1% |
| Unclear | 7 | 21.9% |
| Disagree | 5 | 15.6% |
| Missing | 1 | 3.1% |
| **Total** | 32 | 100% |

Of the 32 uncharacterized glycoproteins, potential annotations, with consistent results from both tools, could be obtained for 3 proteins (Table 5). Their annotations are related to cell-surface functions, which is consistent with the fact that *N*-glycosylation in archaea occurs on the outside of the cell membrane. One protein is annotated as a potential peptide transporter (D4GZ52), the second is likely involved in the S-layer biosynthesis or organization (D4GU84), and the third is annotated to contain a von Willebrand factor domain with potential GATase function (D4GX72) (Supplemental Data 5). For 15 and 1 additional glycoproteins, only Foldseek or eggNOG-mapper returned results, respectively, indicating that structure-based comparisons may be more useful than sequence-based methods for the annotation of uncharacterized proteins in archaea (Table 5). The lower percentage of potential annotations for uncharacterized proteins than for the *P. aeruginosa* dataset is likely related to the higher ratio of proteins of unknown function in archaeal proteomes, highlighting the need for increased experimental, functional characterizations of archaeal proteins. Nevertheless, the ability of ProtPen to functionally annotate several archaeal proteins that are *N*-glycosylated, a modification that has been shown to be involved in various cellular processes, provides valuable insights into the potential roles of glycoproteins in archaea^28,33–36^.

### Limitations

Despite the promising results obtained with ProtPen, several limitations should be acknowledged. The pipeline performance is limited by the coverage and quality of existing orthology and structural databases, including eggNOG and AlphaFold-derived resources. While these databases are actively maintained, their updates follow different schedules than UniProt, which may lead to slight differences in the respectively employed protein sequences. Users should therefore pay attention to the versions of their provided UniProt identifiers.

Furthermore, the accuracy of predictions for truly uncharacterized proteins cannot be directly assessed without experimental validation. Therefore, ProtPen should be viewed as a hypothesis-generating framework that complements, rather than replaces, experimental characterization of protein function. Additionally, the evaluation of ProtPen results was restricted primarily to *Pseudomonas aeruginosa* proteins and utilized a qualitative scoring system that relied on manual inspection of annotation results. While this approach was suitable for the presented proof of concept, a more automated and quantitative consensus mechanism would be desired for large scale applications. The use of large language models to compare the functional descriptors of different tools, or the mapping of protein domains to a standardized gene ontology, e.g. through InterPro2GO^37^, could be explored to achieve this goal.

### Future Directions

Beyond its immediate use for protein annotation, ProtPen’s modular design positions it as a flexible platform for future research and collaborative development. The integration of additional tools is envisioned, such as DeepFri^20^ and Baktfold^38^. Both tools leverage the protein language model ProstT5^39^ to generate Foldseek-compatible structural encodings (3Di sequences) from sequence alone, enabling structure-informed annotation without prior structure prediction. The recently introduced Baktfold also accepts input from Bakta^40^, Prokka^41^, or generic GenBank files, and supports pre-computed structures and custom search databases, giving it considerable flexibility within genome annotation workflows. These tools could serve as alternative or complementary approaches to the Foldseek workflow that is currently implemented in ProtPen, and which relies on AlphaFold structure predictions.

The integration of multiple tools in a unified framework would also facilitate the systematic benchmarking of annotation workflows. The systematic comparison of different tools could not only include general annotation rates for taxonomically diverse datasets, but could also investigate disagreements between tools in more detail. While we have found several instances of disagreement between Foldseek and eggNOG-mapper, comparisons at a larger scale would be necessary to identify patterns or protein properties that result in conflicting annotations, and to determine strategies for how to resolve them. To improve the accessibility of ProtPen for different HPC environments, the current SLURM-based wrappers could also be adapted to other widely used workflow management systems such as Nextflow or Snakemake. We therefore encourage the community to contribute additional tools and workflows, and envision ProtPen to evolve into a shared framework for protein annotation methods.

## Conclusion

ProtPen provides a reproducible and scalable framework for protein function annotation that brings sequence and structure-based approaches together. By combining orthology-based predictions from eggNOG-mapper with structural similarity searches using Foldseek, the pipeline enhances the interpretability and reliability of protein annotations, especially when prioritizing consensus annotation between both tools. The application of ProtPen to biological datasets of bacterial and archaeal species demonstrated its effectiveness in assigning putative functions to uncharacterized proteins that have previously been shown to be of biological importance. Beyond this proof of concept, ProtPen’s modular design and compatibility with different computational platforms (including high-performance computational clusters) make it broadly applicable to diverse organisms and datasets. As new databases and predictive models continue to emerge, the pipeline can be readily extended to incorporate additional tools, supporting future efforts to systematically characterize the functional landscape of proteomes.

## Acknowledgements

We would like to thank Dr. Paul Craig for fruitful discussions and constructive feedback, as well as Braelyn Mcfarland for designing the ProtPen logo and table of contents graphic. We also thank Research Computing at the Rochester Institute of Technology for providing the computational resources and technical support that contributed to the results of this research. This work was supported by the National Institute Of General Medical Sciences of the National Institutes of Health under Award Number R16GM159790. The content of this publication is solely the responsibility of the authors and does not necessarily represent the official views of the National Institutes of Health.

## Supporting Information

**Supplemental Data 1.** ProtPen annotation results for the control dataset, including manual categorizations reflecting agreement between predicted and known protein functions (eggNOG and Foldseek agreement columns).

**Supplemental Data 2.** ProtPen results of the four analyzed prokaryotic reference proteomes, filtered to include only uncharacterized proteins, with separate tabs to indicate our assigned tool performance score.

**Supplemental Data 3.** ProtPen annotation results for uncharacterized proteins from the *Pseudomonas aeruginosa* PAO1 reference proteome for which both Foldseek and eggNOG-mapper returned informative annotations, along with manual categorizations assessing agreement between the two tools (Tool Agreement Assessment column).

**Supplemental Data 4.** ProtPen annotation results for differentially abundant proteins in dataset PXD044807, including manual categorizations reflecting agreement between predicted and known protein functions and assessments of the agreement between the two tools.

**Supplemental Data 5.** ProtPen annotation results for *Haloferax volcanii* glycoproteins, including manual categorizations reflecting agreement between predicted and known protein functions and assessment of the agreement between the two tools.icted and known protein functions and assessment of the agreement between the two tools.

## References

(1) Almeida, A.; Nayfach, S.; Boland, M.; Strozzi, F.; Beracochea, M.; Shi, Z. J.; Pollard, K. S.; Sakharova, E.; Parks, D. H.; Hugenholtz, P.; Segata, N.; Kyrpides, N. C.; Finn, R. D. A Unified Catalog of 204,938 Reference Genomes from the Human Gut Microbiome. Nat Biotechnol 2021, 39 (1), 105–114. 10.1038/s41587-020-0603-3.

(2) Kitts, P. A.; Church, D. M.; Thibaud-Nissen, F.; Choi, J.; Hem, V.; Sapojnikov, V.; Smith, R. G.; Tatusova, T.; Xiang, C.; Zherikov, A.; DiCuccio, M.; Murphy, T. D.; Pruitt, K. D.; Kimchi, A. Assembly: A Resource for Assembled Genomes at NCBI. Nucleic Acids Res 2016, 44 (D1), D73–D80. 10.1093/nar/gkv1226.

(3) Haft, D. H.; DiCuccio, M.; Badretdin, A.; Brover, V.; Chetvernin, V.; O’Neill, K.; Li, W.; Chitsaz, F.; Derbyshire, M. K.; Gonzales, N. R.; Gwadz, M.; Lu, F.; Marchler, G. H.; Song, J. S.; Thanki, N.; Yamashita, R. A.; Zheng, C.; Thibaud-Nissen, F.; Geer, L. Y.; Marchler-Bauer, A.; Pruitt, K. D. RefSeq: An Update on Prokaryotic Genome Annotation and Curation. Nucleic Acids Res 2018, 46 (D1), D851–D860. 10.1093/nar/gkx1068.

(4) Goodacre, N. F.; Gerloff, D. L.; Uetz, P. Protein Domains of Unknown Function Are Essential in Bacteria. mBio 2013, 5 (1), 10.1128/mbio.00744-13. 10.1128/mbio.00744-13.

(5) Hartman, A. L.; Norais, C.; Badger, J. H.; Delmas, S.; Haldenby, S.; Madupu, R.; Robinson, J.; Khouri, H.; Ren, Q.; Lowe, T. M.; Maupin-Furlow, J.; Pohlschroder, M.; Daniels, C.; Pfeiffer, F.; Allers, T.; Eisen, J. A. The Complete Genome Sequence of Haloferax Volcanii DS2, a Model Archaeon. PLoS One 2010, 5 (3), e9605. 10.1371/journal.pone.0009605.

(6) Pfeiffer, F.; Oesterhelt, D.; Pfeiffer, F.; Oesterhelt, D. A Manual Curation Strategy to Improve Genome Annotation: Application to a Set of Haloarchael Genomes. Life 2015, 5 (2), 1427–1444. 10.3390/life5021427.

(7) Stover, C. K.; Pham, X. Q.; Erwin, A. L.; Mizoguchi, S. D.; Warrener, P.; Hickey, M. J.; Brinkman, F. S. L.; Hufnagle, W. O.; Kowalik, D. J.; Lagrou, M.; Garber, R. L.; Goltry, L.; Tolentino, E.; Westbrock-Wadman, S.; Yuan, Y.; Brody, L. L.; Coulter, S. N.; Folger, K. R.; Kas, A.; Larbig, K.; Lim, R.; Smith, K.; Spencer, D.; Wong, G. K.-S.; Wu, Z.; Paulsen, I. T.; Reizer, J.; Saier, M. H.; Hancock, R. E. W.; Lory, S.; Olson, M. V. Complete Genome Sequence of Pseudomonas Aeruginosa PAO1, an Opportunistic Pathogen. Nature 2000, 406 (6799), 959–964. 10.1038/35023079.

(8) Camacho, C.; Coulouris, G.; Avagyan, V.; Ma, N.; Papadopoulos, J.; Bealer, K.; Madden, T. L. BLAST+: Architecture and Applications. BMC Bioinformatics 2009, 10 (1), 421. 10.1186/1471-2105-10-421.

(9) Potter, S. C.; Luciani, A.; Eddy, S. R.; Park, Y.; Lopez, R.; Finn, R. D. HMMER Web Server: 2018 Update. Nucleic Acids Res 2018, 46 (W1), W200–W204. 10.1093/nar/gky448.

(10) Cantalapiedra, C. P.; Hernández-Plaza, A.; Letunic, I.; Bork, P.; Huerta-Cepas, J. eggNOG-Mapper v2: Functional Annotation, Orthology Assignments, and Domain Prediction at the Metagenomic Scale. Mol Biol Evol 2021, 38 (12), 5825–5829. 10.1093/molbev/msab293.

(11) Buchfink, B.; Xie, C.; Huson, D. H. Fast and Sensitive Protein Alignment Using DIAMOND. Nat Methods 2015, 12 (1), 59–60. 10.1038/nmeth.3176.

(12) Steinegger, M.; Söding, J. MMseqs2 Enables Sensitive Protein Sequence Searching for the Analysis of Massive Data Sets. Nat Biotechnol 2017, 35 (11), 1026–1028. 10.1038/nbt.3988.

(13) Huerta-Cepas, J.; Szklarczyk, D.; Heller, D.; Hernández-Plaza, A.; Forslund, S. K.; Cook, H.; Mende, D. R.; Letunic, I.; Rattei, T.; Jensen, L. J.; von Mering, C.; Bork, P. eggNOG 5.0: A Hierarchical, Functionally and Phylogenetically Annotated Orthology Resource Based on 5090 Organisms and 2502 Viruses. Nucleic Acids Res 2019, 47 (D1), D309–D314. 10.1093/nar/gky1085.

(14) Kulmanov, M.; Khan, M. A.; Hoehndorf, R. DeepGO: Predicting Protein Functions from Sequence and Interactions Using a Deep Ontology-Aware Classifier. Bioinformatics 2018, 34 (4), 660–668. 10.1093/bioinformatics/btx624.

(15) Ardern, Z.; Chakraborty, S.; Lenk, F.; Kaster, A.-K. Elucidating the Functional Roles of Prokaryotic Proteins Using Big Data and Artificial Intelligence. FEMS Microbiol Rev 2023, 47 (1), fuad003. 10.1093/femsre/fuad003.

(16) Jumper, J.; Evans, R.; Pritzel, A.; Green, T.; Figurnov, M.; Ronneberger, O.; Tunyasuvunakool, K.; Bates, R.; Žídek, A.; Potapenko, A.; Bridgland, A.; Meyer, C.; Kohl, S. A. A.; Ballard, A. J.; Cowie, A.; Romera-Paredes, B.; Nikolov, S.; Jain, R.; Adler, J.; Back, T.; Petersen, S.; Reiman, D.; Clancy, E.; Zielinski, M.; Steinegger, M.; Pacholska, M.; Berghammer, T.; Bodenstein, S.; Silver, D.; Vinyals, O.; Senior, A. W.; Kavukcuoglu, K.; Kohli, P.; Hassabis, D. Highly Accurate Protein Structure Prediction with AlphaFold. Nature 2021, 596 (7873), 583–589. 10.1038/s41586-021-03819-2.

(17) Berman, H. M.; Westbrook, J.; Feng, Z.; Gilliland, G.; Bhat, T. N.; Weissig, H.; Shindyalov, I. N.; Bourne, P. E. The Protein Data Bank. Nucleic Acids Res 2000, 28 (1), 235–242. 10.1093/nar/28.1.235.

(18) Varadi, M.; Bertoni, D.; Magana, P.; Paramval, U.; Pidruchna, I.; Radhakrishnan, M.; Tsenkov, M.; Nair, S.; Mirdita, M.; Yeo, J.; Kovalevskiy, O.; Tunyasuvunakool, K.; Laydon, A.; Žídek, A.; Tomlinson, H.; Hariharan, D.; Abrahamson, J.; Green, T.; Jumper, J.; Birney, E.; Steinegger, M.; Hassabis, D.; Velankar, S. AlphaFold Protein Structure Database in 2024: Providing Structure Coverage for over 214 Million Protein Sequences. Nucleic Acids Res 2024, 52 (D1), D368–D375. 10.1093/nar/gkad1011.

(19) Holm, L.; Laiho, A.; Törönen, P.; Salgado, M. DALI Shines a Light on Remote Homologs: One Hundred Discoveries. Protein Science 2023, 32 (1), e4519. 10.1002/pro.4519.

(20) Gligorijević, V.; Renfrew, P. D.; Kosciolek, T.; Leman, J. K.; Berenberg, D.; Vatanen, T.; Chandler, C.; Taylor, B. C.; Fisk, I. M.; Vlamakis, H.; Xavier, R. J.; Knight, R.; Cho, K.; Bonneau, R. Structure-Based Protein Function Prediction Using Graph Convolutional Networks. Nat Commun 2021, 12 (1), 3168. 10.1038/s41467-021-23303-9.

(21) van Kempen, M.; Kim, S. S.; Tumescheit, C.; Mirdita, M.; Lee, J.; Gilchrist, C. L. M.; Söding, J.; Steinegger, M. Fast and Accurate Protein Structure Search with Foldseek. Nat Biotechnol 2024, 42 (2), 243–246. 10.1038/s41587-023-01773-0.

(22) Ruperti, F.; Papadopoulos, N.; Musser, J. M.; Mirdita, M.; Steinegger, M.; Arendt, D. Cross-Phyla Protein Annotation by Structural Prediction and Alignment. Genome Biol 2023, 24, 113. 10.1186/s13059-023-02942-9.

(23) Bouras, G.; Grigson, S. R.; Mirdita, M.; Heinzinger, M.; Papudeshi, B.; Mallawaarachchi, V.; Green, R.; Kim, R. S.; Mihalia, V.; Psaltis, A. J.; Wormald, P.-J.; Vreugde, S.; Steinegger, M.; Edwards, R. A. Protein Structure-Informed Bacteriophage Genome Annotation with Phold.

(24) Heinzinger, M.; Weissenow, K.; Sanchez, J. G.; Henkel, A.; Mirdita, M.; Steinegger, M.; Rost, B. Bilingual Language Model for Protein Sequence and Structure. NAR Genom Bioinform 2024, 6 (4), lqae150. 10.1093/nargab/lqae150.

(25) Queirós, P.; Delogu, F.; Hickl, O.; May, P.; Wilmes, P. Mantis: Flexible and Consensus-Driven Genome Annotation. GigaScience 2021, 10 (6), giab042. 10.1093/gigascience/giab042.

(26) UniProt Consortium. UniProt: The Universal Protein Knowledgebase in 2021. Nucleic Acids Res 2021, 49 (D1), D480–D489. 10.1093/nar/gkaa1100.

(27) Wang, M.; Medarametla, P.; Kronenberger, T.; Deingruber, T.; Brear, P.; Figueroa, W.; Ho, P.-M.; Krueger, T.; Pearce, J. C.; Poso, A.; Wakefield, J. G.; Spring, D. R.; Welch, M. Pseudomonas Aeruginosa Acyl-CoA Dehydrogenases and Structure-Guided Inversion of Their Substrate Specificity. Nat Commun 2025, 16 (1), 2334. 10.1038/s41467-025-57532-z.

(28) Schulze, S.; Pfeiffer, F.; Garcia, B. A.; Pohlschroder, M. Comprehensive Glycoproteomics Shines New Light on the Complexity and Extent of Glycosylation in Archaea. PLOS Biology 2021, 19 (6), e3001277. 10.1371/journal.pbio.3001277.

(29) Alhusseini, L. B.; Hasani, B.; Jaafar, F. N.; Beig, M.; Abbasian, S.; Azizian, K. Temporal and Geographic Trends in Extended-Spectrum Cephalosporins Resistance among Neisseria Gonorrhoeae Isolates Worldwide: A Systematic Review and Meta-Analysis. BMC Infect Dis 2025, 25 (1), 1175. 10.1186/s12879-025-11601-2.

(30) Wood, S. J.; Kuzel, T. M.; Shafikhani, S. H. Pseudomonas Aeruginosa: Infections, Animal Modeling, and Therapeutics. Cells 2023, 12 (1), 199. 10.3390/cells12010199.

(31) Pohlschroder, M.; Schulze, S.; Pfeiffer, F.; Hong, Y. Haloferax Volcanii: A Versatile Model for Studying Archaeal Biology. Journal of Bacteriology 2025, 207 (6), e00062–25. 10.1128/jb.00062-25.

(32) Kengen, S. W. M. ‘Pyrococcus Furiosus, 30 Years On.’ Microbial Biotechnology 2017, 10 (6), 1441–1444. 10.1111/1751-7915.12695.

(33) Shalev, Y.; Turgeman-Grott, I.; Tamir, A.; Eichler, J.; Gophna, U. Cell Surface Glycosylation Is Required for Efficient Mating of Haloferax Volcanii. Front Microbiol 2017, 8, 1253. 10.3389/fmicb.2017.01253.

(34) Esquivel, R. N.; Schulze, S.; Xu, R.; Hippler, M.; Pohlschroder, M. Identification of Haloferax Volcanii Pilin N-Glycans with Diverse Roles in Pilus Biosynthesis, Adhesion, and Microcolony Formation. J Biol Chem 2016, 291 (20), 10602–10614. 10.1074/jbc.M115.693556.

(35) Tripepi, M.; You, J.; Temel, S.; Önder, Ö.; Brisson, D.; Pohlschröder, M. N-Glycosylation of Haloferax Volcanii Flagellins Requires Known Agl Proteins and Is Essential for Biosynthesis of Stable Flagella. J Bacteriol 2012, 194 (18), 4876–4887. 10.1128/JB.00731-12.

(36) Tamir, A.; Eichler, J. N-Glycosylation Is Important for Proper Haloferax Volcanii S-Layer Stability and Function. Appl Environ Microbiol 2017, 83 (6), e03152–16. 10.1128/AEM.03152-16.

(37) Mitchell, A.; Chang, H.-Y.; Daugherty, L.; Fraser, M.; Hunter, S.; Lopez, R.; McAnulla, C.; McMenamin, C.; Nuka, G.; Pesseat, S.; Sangrador-Vegas, A.; Scheremetjew, M.; Rato, C.; Yong, S.-Y.; Bateman, A.; Punta, M.; Attwood, T. K.; Sigrist, C. J. A.; Redaschi, N.; Rivoire, C.; Xenarios, I.; Kahn, D.; Guyot, D.; Bork, P.; Letunic, I.; Gough, J.; Oates, M.; Haft, D.; Huang, H.; Natale, D. A.; Wu, C. H.; Orengo, C.; Sillitoe, I.; Mi, H.; Thomas, P. D.; Finn, R. D. The InterPro Protein Families Database: The Classification Resource after 15 Years. Nucleic Acids Res 2015, 43 (D1), D213–D221. 10.1093/nar/gku1243.

(38) Bouras, G.; Lim, S. won; Durr, L.; Vreugde, S.; Goesmann, A.; Edwards, R. A.; Schwengers, O. Baktfold: Sensitive Protein Functional Annotation across the Microbial Tree of Life Using Structural Information. bioRxiv April 1, 2026, p 2026.03.31.715528. 10.64898/2026.03.31.715528.

(39) Heinzinger, M.; Weissenow, K.; Sanchez, J. G.; Henkel, A.; Steinegger, M.; Rost, B. ProstT5: Bilingual Language Model for Protein Sequence and Structure. bioRxiv July 25, 2023, p 2023.07.23.550085. 10.1101/2023.07.23.550085.

(40) Schwengers, O.; Jelonek, L.; Dieckmann, M. A.; Beyvers, S.; Blom, J.; Goesmann, A. Bakta: Rapid and Standardized Annotation of Bacterial Genomes via Alignment-Free Sequence Identification: Find out More about Bakta, the Motivation, Challenges and Applications, Here. Microbial Genomics 2021, 7 (11). 10.1099/mgen.0.000685.

(41) Seemann, T. Prokka: Rapid Prokaryotic Genome Annotation. Bioinformatics 2014, 30 (14), 2068–2069. 10.1093/bioinformatics/btu153.

